# A Cesium Chloride Gradient Ultracentrifugation-Based Method for the Isolation of DNA from Diverse Recalcitrant Plant Species for Nanopore Sequencing

**DOI:** 10.64898/2026.08.18.745475

**Authors:** June Labbancz, Amit Dhingra

**Affiliations:** Department of Horticultural Sciences, Texas A&M University, College Station, Texas 77843, USA

**Keywords:** DNA isolation, Long-read sequencing, Plant Genomics

## Abstract

Developments in Nanopore sequencing have enabled telomere to telomere genomic assembly as a routine technique in genomic research. Nanopore DNA sequencing for genomic assembly is typically performed on native DNA molecules, making it particularly sensitive to the quality of input DNA, with contaminating molecules limiting data yields and reducing read quality. As pangenome analysis gains interest, particularly in non-model plant species which are often rich in inhibitory secondary metabolites, the development of methods which can improve the quality and throughput of nanopore sequencing is essential. Here we describe a method for isolation of total DNA from the leaf tissues of diverse Viridiplantae species. The initial lysis buffer consists of a modified CTAB buffer, incorporating dimethyl sulfoxide for the reduction of viscosity, which can be problematic in many plant DNA preparations. An organic extraction with 2-butoxyethanol is utilized to further extract phenolic compounds which may be sufficiently hydrophilic to evade chloroform extraction, while reducing aqueous phase volume. Further cleanup via cesium chloride (CsCl) ultracentrifugation is performed to minimize the carryover of residual contaminating macromolecules. Samples prepared using this method are of consistent high quality, even when extracted from challenging late season leaf tissue or secondary metabolite rich species. Sequencing results from samples prepared by this method outperform those obtained from typical modified CTAB DNA isolation techniques in both quantity and quality. We tested sequencing performance from *Vitis* DNA isolated using a modified CTAB method and *Vitis* DNA isolated using the CsCl ultracentrifugation-based method described here. DNA isolated via the method described here produced 83% more >Q10 sequence data (52.61 Gb vs. 28.8 Gb), resulted in a 60% greater read N50 despite more handling steps (32.78kb vs. 20.45kb), and resulted in a higher modal read quality (Q27 vs. Q24). The consistency of this method across diverse plant taxa suggests its use as a general method for DNA isolation prior to Nanopore sequencing and genomic assembly for diverse plant taxa.

## Background

Advancements in Nanopore sequencing have made high-quality genomic assembly increasingly attainable, with single platform telomere to telomere assembly now possible [1]. These advancements have ushered in a new era of pangenome analysis, where collections of dozens to hundreds of genomes may be sequenced and assembled to discover trait-linked genetic diversity and core genetic architecture across species and even genera [2,3]. Plant species which have previously had no published genomes, including less cultivated horticultural crops, are also being sequenced and assembled for the first time [4]. Difficulties in isolating DNA of acceptable quality for sequencing and genomic assembly have been identified as a limitation in many plant lineages, with woody plants often being identified as particularly problematic [5]. Even within a genus, variation may exist in the ease of DNA isolation [6,7]. To facilitate the study of a wider sampling of plant genetic diversity, there is a clear need for improved methods to isolate DNA of acceptable quality for whole genome sequencing.

The most challenging contaminants in DNA isolations from plant species are typically polysaccharides and phenolic compounds [5,8]. As these molecules can become entangled with or bound to DNA or possess physical properties which are similar enough to that of DNA to avoid complete removal, they often remain in DNA preparations for downstream processes, such as sequencing, resulting in inferior experimental results [8]. Polymerase Chain Reaction (PCR) amplification is a typical step in the preparation of sequencing libraries for some sequencing platforms (such as in Illumina sequencing). This can increase the total quantity of DNA as well as the quantity of DNA relative to unamplified contaminants provided the reaction is not fully inhibited, effectively increasing sample quality. Sequencing of native DNA molecules, however, can be particularly advantageous as it reduces bias, enables longer sequencing reads, and enables the sequencing of epigenetic modifications [9,10]. As such, the need for greater quantities of high-quality DNA for sequencing has increased.

The most common methods for the isolation of DNA in plant science are silica column-based kits and the modified CTAB protocol [11,12]. The CTAB-based DNA isolation most famously described by Doyle and Doyle is regularly modified, although modifications are typically minor, including variations in incubation time, beta-mercaptoethanol concentration, and the addition of polyvinylpyrrolidone [12,13]. Less frequently used methods include nuclei isolation-based methods, methods incorporating enzymatic polysaccharide removal, and alternative detergent systems, but often still face taxon-specific issues [14–16]. Cesium chloride ultracentrifugation is a well-described method for the isolation of DNA, generally yielding high-quality DNA relative to CTAB-based methods, although it is readily identified as a method which has fallen into disuse [17–19]. A CsCl ultracentrifugation-based protocol for DNA isolation from the chlorophyte *Prototheca wickerhamii* for NGS sequencing has been previously described [20], but a broadly applicable method for sequencing-ready DNA from vascular plants remains undescribed. Some macromolecules may still co-band with DNA in a CsCl gradient, including some polysaccharides [21,22]. This makes a protocol which incorporates complimentary lysis and cleanup steps in addition to CsCl-ultracentrifugation essential for experimental success in the face of plant taxa which contain diverse and heterogenous mixtures of polysaccharides and phenolic compounds.

To confront the challenge of isolating sequencing-quality DNA from diverse recalcitrant plant taxa, we developed a method which incorporates a modified CTAB buffer including DMSO for lysis and organic extraction, 2-butoxyethanol salt-assisted liquid-liquid extraction to further remove phenolic compounds and reduce aqueous volume while avoiding precipitation, followed by CsCl ultracentrifugation. Following ultracentrifugation, a further 2-butoxyethanol salt-assisted liquid-liquid extraction at low pH is carried out, followed by PEG precipitation and repeated ethanol washes. To evaluate the broad applicability of the method described here, we compared DNA isolations of 17 samples from across Viridiplantae (Figure 1) by 3 methods: a silica column-based method (DNEasy Plant Pro Kit), a modified CTAB method, and a novel CsCl-based method. Three *Vitis* samples were used to generate a DNA sequencing library using a modified CTAB method and a novel CsCl-based method. Data generated in one sequencing run on an Oxford Nanopore R10.4.1 flow cell was used to compare the effect of the novel method on sequencing output.

**Figure 1.**
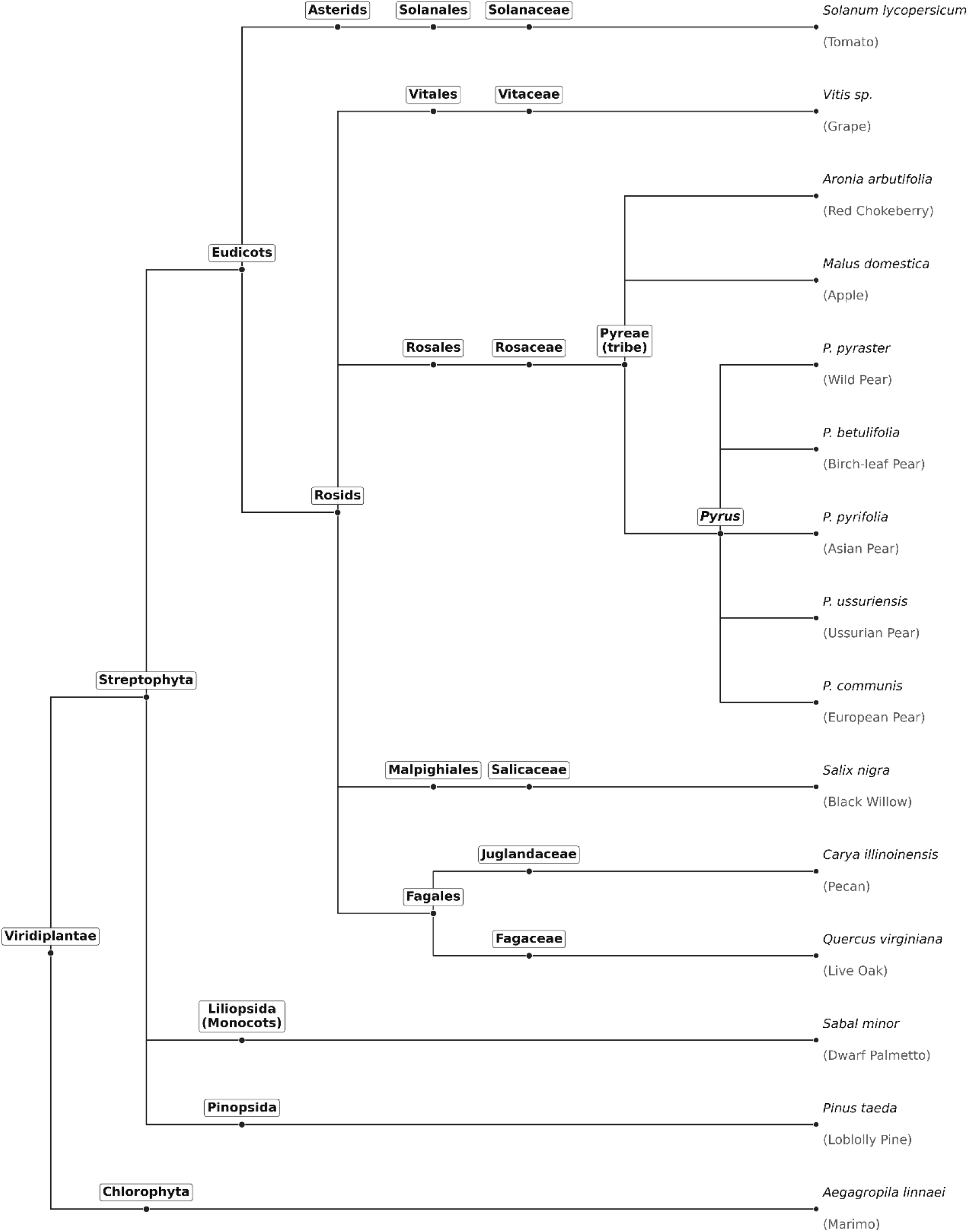
Taxonomic relationships of organisms sampled in this study.

## Methods

### Sample Collection and Preparation

All leaf samples were placed in a 50mL polypropylene conical tube and flash frozen in liquid nitrogen prior to milling. *Pyrus pyrifolia, Pyrus ussuriensis*, and *Pyrus betulifolia* leaf samples were held at 4°C overnight for shipment prior to flash freezing in liquid nitrogen. Samples were cryogenically milled using a Spex Sample/Prep FreezerMill 6875 for 3 minutes at 15 cpm, then stored at -80°C until use.

### Buffer Preparations

Isolation Buffer: 3% CTAB, 1% Sarkosyl, 1.7M NaCl, 100mM LiCl, 100mM Tris-HCl pH8, 25mM EDTA pH8

CsCl Buffer (1.86g/mL): Add 180g CsCl to 100mL Buffer TE, mix well. Incubate at 42°C with intermittent agitation until fully dissolved.

CsCl Buffer (1.56g/mL): Add 104g CsCl to 100mL Buffer TE, mix well. Incubate at 42°C with intermittent agitation until fully dissolved.

CsCl Loading Buffer: Add 10µL β-mercaptoethanol per 1mL and mix gently. Add 1µL SYBR Gold per 1mL and mix gently. Prepare only immediately before loading.

Citrate Buffer: 100mM Citric Acid, 1M NaCl. Adjust to pH 3.5 with NaOH.

PEG Precipitation Buffer: 12.5% PEG8000, 1M NaCl, 100mM Tris-HCl pH8, 25mM EDTA pH8

### CsCl Ultracentrifugation Method

1. Add 1% polyvinylpolypyrrolidone to 4mL of Isolation Buffer in a 15mL Lo-Bind DNA tube and heat to 60°C in a water bath.
2. Add 450µL dimethyl sulfoxide and 200µL β-mercaptoethanol to the tube, gently inverting between additions.
3. Add 300mg of tissue per tube, close the tube, then shake well.
4. Incubate at 60°C for 30 minutes, regularly agitating.
5. Add 7mL 24:1 chloroform: isoamylalcohol, mix well by hand. Put on a rotational mixer for 5 minutes.
6. Centrifuge at 3000g for 10 minutes at room temperature.
7. Transfer 3mL supernatant to a fresh 5mL tube. Add 2mL 2-butoxyethanol, mix well by hand.
8. Centrifuge at 10000g for 5 minutes at room temperature.
9. Remove the aqueous (bottom) phase by pipette to a fresh 5mL tube, avoiding precipitate if any is present.
10. Add 2.5mL 2-butoxyethanol, mix well by hand.
11. Centrifuge again at 10000g for 5 minutes at room temperature.
12. Transfer aqueous (bottom) phase by pipette to a fresh 5mL tube.
13. Add sterile water to the solution until it reaches 2.1mL volume.
14. Add 3mL 1.86g/mL CsCl solution 1mL at a time, mixing between additions.
15. Add 500µL CsCl loading buffer, then gently mix.
16. Load sample into 5PP seal tubes using <18-gauge needles, weigh all tubes to ensure balancing (<100mg difference), and seal all tubes. Insert into vertical centrifuge rotor (P100VT in this study).
17. Centrifuge at 83,000g for 20-24 hours.
18. Unload all tubes for the ultracentrifuge rotor and make a puncture using a fresh needle at the top of the tube.
19. Visualize the DNA band using a 450-500nm flashlight. Using a <18-gauge needle and 1mL syringe, make a puncture ∼2mm below the band, then extract the DNA. Ideally extract the DNA in less than 0.5mL solution. Transfer to a 5mL tube.
20. Add 1mL Citrate Buffer, mix gently, then add 2mL 2-butoxyethanol.
21. Centrifuge at 10000g for 3 minutes.
22. Transfer aqueous phase (bottom) to a fresh 5mL tube and immediately add 4mL PEG Precipitation Buffer. (Note: it is imperative that time spent at low pH from step 20 to 22 is minimized).
23. Incubate at 4ºC for 1 hour.
24. Centrifuge at 12,000g for 20 minutes.
25. Carefully remove the supernatant by pipette, as the pellet may not be visible.
26. Add 2mL 70% ethanol, invert gently. Centrifuge at 12,000g for 3 minutes.
27. Remove supernatant and repeat Step 26.
28. Remove supernatant and resuspend in 100µL Buffer TE (10mM Tris-HCl pH 8, 1mM EDTA pH 8). Incubate at 37°C for 5 minutes.

### Comparison Methods

A modified CTAB protocol was followed as described. 1mL of CTAB Isolation Buffer (3% CTAB, 1.4M NaCl, 100mM Tris-HCl pH8, 50mM EDTA pH 8.0) was heated to 60°C, add 2.5% β-mercaptoethanol immediately prior to the addition of sample. 100mg of milled sample was added to the buffer and immediately mixed well by hand. Samples were incubated at 60°C for 30 minutes with regular agitation prior to the addition of an equal volume of 24:1 chloroform: isoamylalcohol. The samples were centrifuged at 8,000g for 6 minutes at room temperature. The supernatant was moved to a fresh tube, and the chloroform: isoamylalcohol extraction repeated. An equal volume of isopropanol was added, mixed gently by hand, then incubated at 4°C for 30 minutes. Samples were centrifuged at 12,000g for 12 minutes, then the pellet washed twice with 70% ethanol. The pellet was resuspended in 700µL TE with 4µL/mL RNAse A, then incubated at 42°C for 3 hours to accommodate the long time for the pellet of some samples to fully redissolve. 300µL Proteinase K solution was added (250mM NaCl, 1% sarkosyl, 1% β-mercaptoethanol, 2µL/mL Proteinase K) was added and samples were incubated for 15 minutes. A single chloroform: isoamylalcohol extraction was performed as described above, then the sample was precipitated with equal volume isopropanol, then incubated at 4°C for 30 minutes. Samples were centrifuged at 12,000g for 12 minutes, then the pellet washed twice with 70% ethanol. Samples were resuspended in 100µL Buffer TE.

The DNEasy Plant Pro Kit (Qiagen) was used according to the manufacturer protocol with the deviation of eluting into 2×100µL Buffer TE rather than the manufacturer provided Buffer AE. All optional but recommended steps were followed. 75mg of sample input was used per isolation.

For all samples by all methods, 3 replicates each were tested from the same milled tissue sample. Final eluted or resuspended isolations were quantified on the NanoDrop 8000 UV-Vis spectrophotometer (ThermoFisher Scientific, USA) using Buffer TE as a blank and quantified using a Qubit 4 Fluorometer (ThermoFisher Scientific, USA) with the dsDNA Broad Range Assay.

To assess the removal of pigmented compounds by 2-butoxyethanol extraction, an additional isolation was prepared from two samples, *Carya illinoinensis* cv. ‘Pawnee’ and S*abal minor*. After steps 6 and 9, 1mL of aqueous sample was pipetted into a cuvette and absorbance at 1nm intervals from 750nm to 400nm was measured with a M4 UV/Vis Spectrophotometer (VWR, USA).

### DNA Sequencing

DNA libraries were prepared using samples from three *Vitis* samples ‘Lomanto’, ‘Munson’, and ‘America’. Prior to library preparation, the Short Fragment Eliminator Kit (Oxford Nanopore, UK) was used on all samples according to manufacturer protocol. Samples were resuspended in 20µL Buffer TE then quantified using Qubit fluorometry. 900-1000ng of DNA was used for each sample for library preparation using the Native Barcoding 24 Kit 24 V14 (Oxford Nanopore, UK). The library preparation was completed according to manufacturer protocol, with one alteration in the form of extended incubation times during the elution of DNA from the Ampure XP beads. Final DNA libraries were quantified using the Qubit 4 dsDNA Broad Range Assay.

70-80ng total DNA library was loaded into an R10.4.1 flow cell on a Promethion P2 Solo device, and an experiment was run until the total pore count reached 1700 pores or less.

## Results and Discussion

Spectrophotometric assessment shows near indifference of the CsCl-based method to the sample used, with improvements on samples isolated via silica column and CTAB-based methods in all samples (Table 1). To facilitate comparison between samples, a quality threshold was established: 1) a 260/280 ratio between 1.7 and 1.9, a 260/230 ratio between 2 and 3, and a Nanodrop/Qubit DNA quantity ratio <2. Optimal 260/280 and 260.230 ratios of ∼1.8 and >2, respectively, have been long established for DNA UV spectrophotometry. Assessing DNA purity by comparing the DNA-specific fluorescence measurement of the Qubit with the relatively non-specific A260 can identify contaminants which may have a wide absorption spectrum, or RNA presence due to issues such as inhibition of RNAse during DNA preparation. While exact cutoff criteria are not universally agreed upon, a large discordance between the two measures is a potential flag for poor DNA isolation quality. All but one sample (with a Qubit/Nanodrop ratio of 2.12) passed the established quality threshold using the CsCl-based protocol, with few of the silica column and CTAB-based preparations passing all three quality criteria (Table 2). Four of the samples isolated by the modified CTAB method (*Aronia arbutifolia* and three *Pyrus* samples, all in subtribe Malinae) failed to resuspend following final resuspension after 24 hours of incubation at 37°C, forming a semi-solid gel, preventing quantification and qualification. A failure to resuspend has been identified as a failure mode of DNA isolations previously [23].

**Table 1.**
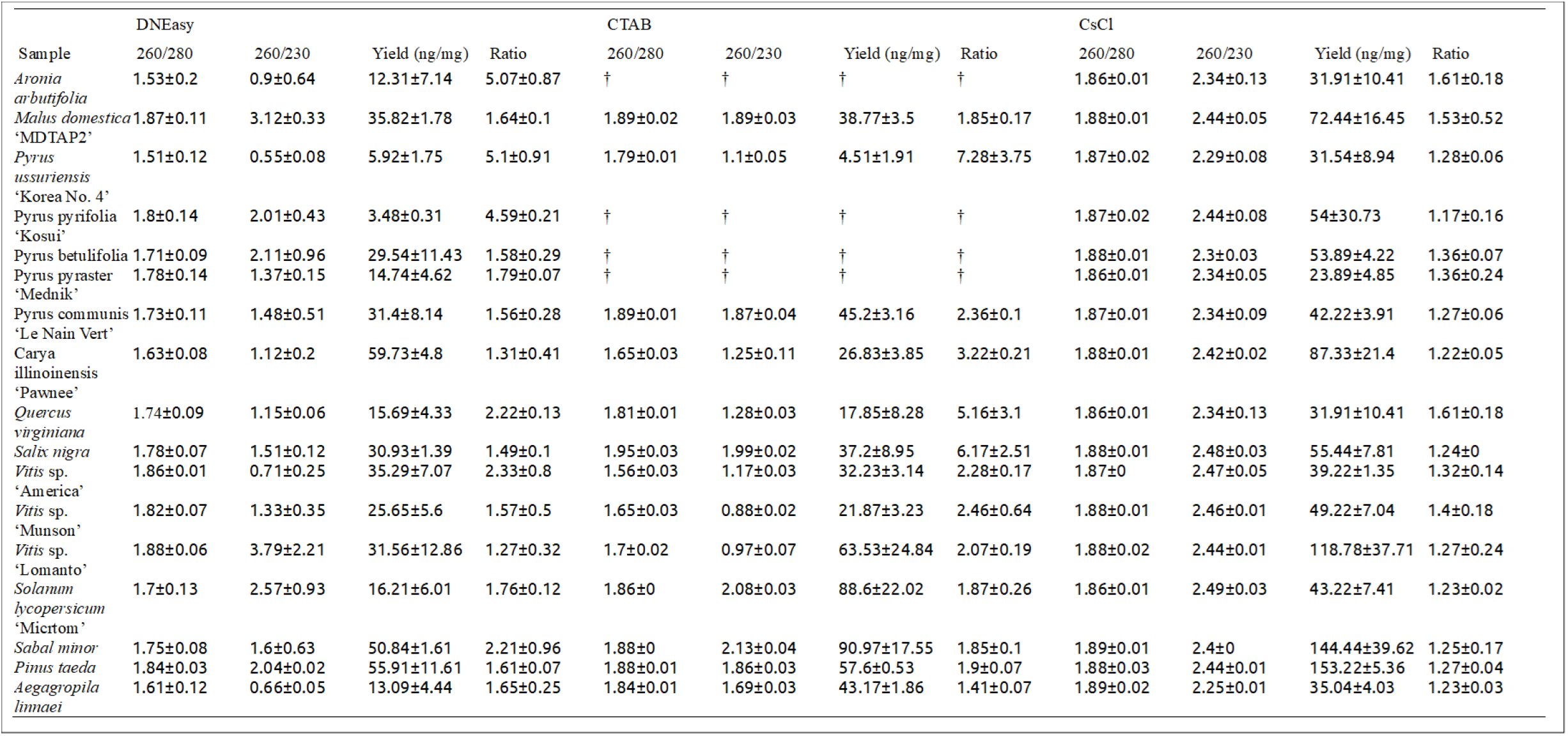
Qualification and quantification of DNA isolated from 17 samples by 3 methods. † Samples which did not resuspend after 24 hours, forming a semi-solid gel.

**Table 2.** Number of samples out of the total (51 - 17 samples x 3 replicates) passing established quality criteria.

| Metric | Silica Column | CTAB | CsCl |
| --- | --- | --- | --- |
| 260/280 | 31/51 | 25/51 | 51/51 |
| 260/230 | 10/51 | 7/51 | 51/51 |
| Nanodrop/Qubit | 36/51 | 15/51 | 50/51 |
| All Three | 9/51 | 5/51 | 50/51 |

Previous DNA isolation methods have identified many species within Malinae as recalcitrant, being rich in polysaccharides such as pectin and polyphenolics such as arbutin [24,25]. The method presented here was able to isolate sequencing-ready DNA from each of the 4 samples which failed to resuspend in the CTAB-based method (Table 1). Ideally, DNA isolation methods would be minimally taxon-specific, minimizing the need for troubleshooting and adaptation of methods between taxa dependent on specific metabolites. By this metric, the CsCl ultracentrifugation-based method is particularly attractive relative to standard methods in use today.

Comparing sequencing outputs from DNA isolated via a modified CTAB method and via the method presented here, major improvements are seen in sequencing yield, read length, and read quality (Table 3, Figure 2). Genome assembly is sensitive to these metrics, with an adequate amount of data, with reads long enough to consistently bridge repetitive regions, with minimal sequencing errors being suggested for optimal results. Coverage less than 50X may result in excessively fragmented plant genome assemblies, for example [26]. Using the 50X coverage recommendation in *Vitis* (25Gb for a ∼500Mb haploid genome size), for example, the CsCl-based method would be adequate for two genomic assemblies, while the CTAB-based method would only be adequate for one. Read N50 was lower in the sequencing run on DNA isolated via the modified CTAB method, despite the utilization of two size selection methods for both samples. Even relatively small changes in read N50 can result in large differences in assembly contiguity; in maize at 50X coverage, an increase in N50 from 16kb to 21kb resulted in an increase in contig N50 from about 4Mb to 16Mb [26]. Overall, the DNA yielded from the CsCl ultracentrifugation-based method generated data more amenable to genomic assembly, which is dependent upon adequate sequence data quantity, long sequencing reads, and reads with minimal errors present.

**Table 3.** Summary statistics for a single sequencing run on three *Vitis* samples by two methods. Percent difference in read quality/error rate calculated by applying the the formula P=10^-Q/10^ to the observed data for passed reads: 10^-22.3/10^/10^-26.4/10^.

| Attribute | CTAB Method | CsCl Method | Difference |
| --- | --- | --- | --- |
| Total Generated Data | 31.3 Gb | 53.9 Gb | +72.2% |
| Data Passing Quality Threshold (>Q10) | 28.8 Gb | 52.6 Gb | +82.6% |
| Read N50 | 20.45 kb | 32.78 kb | +60.2% |
| Median Read Quality (Passed Reads) | 22.3 | 26.4 | +257% |

**Figure 2.**
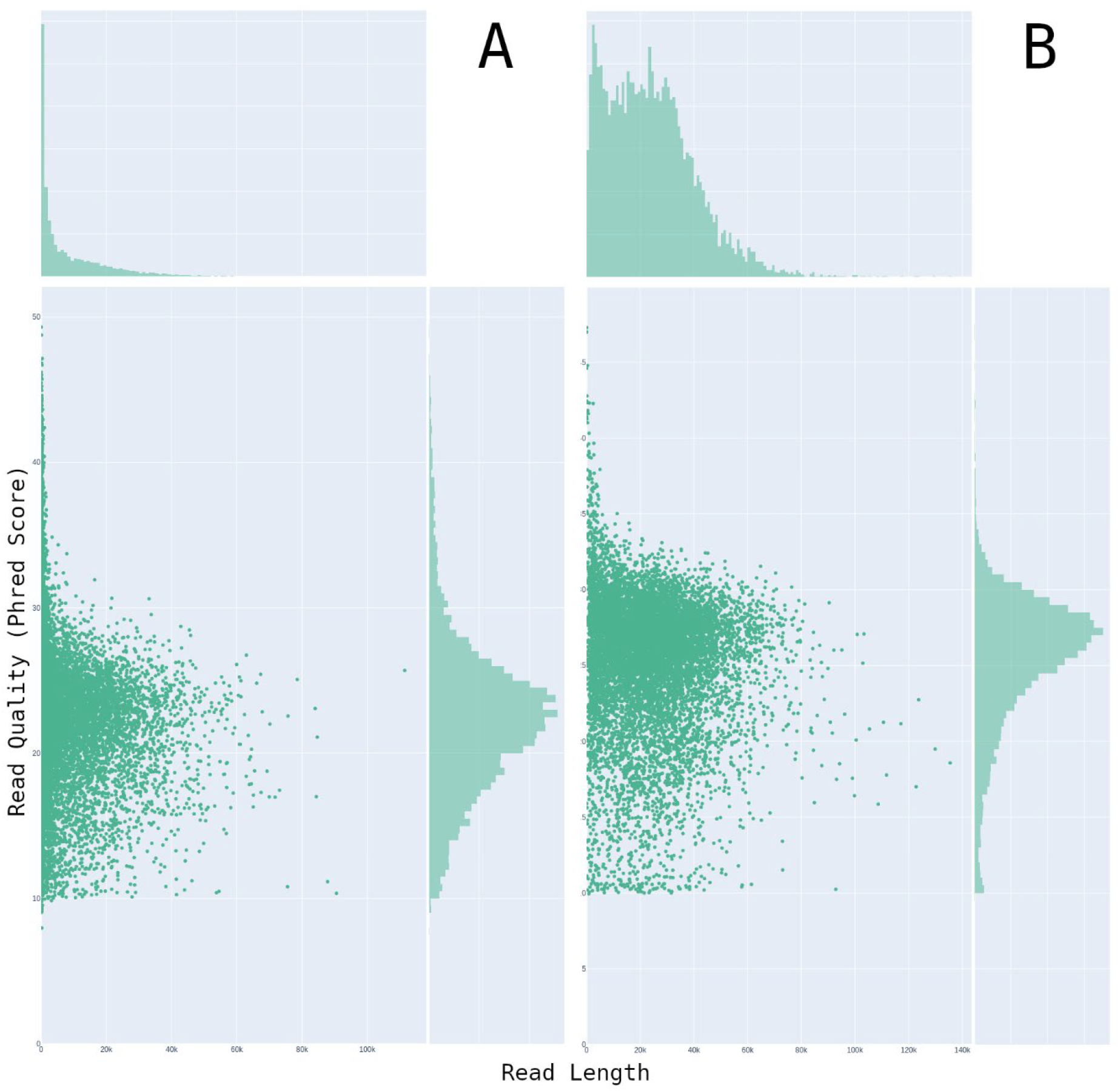
Plot of passed reads by read quality and read length in sequencing experiments using samples from A) a modified CTAB method and B) the method presented in this manuscript. The x-axis scale is different between A and B.

Although 2-butoxyethanol has been previously used as a reagent to sequentially precipitate contaminants then DNA [27], a liquid-liquid extraction utilizing 2-butoxyethanol for DNA purification has not been previously described. 2-butoxyethanol is known to exhibit phase separation with water under certain conditions, such as elevated temperature and salt content [28,29]. This behavior can be exploited to use it for organic extraction. The greater hydrophilicity of 2-butoxyethanol relative to the commonly used phenol and chloroform (logP of 0.83 vs. 1.5 and 1.97, respectively) may allow it to extract a greater range of compounds such as plant phenolic compounds [30,31]. Using visible wavelength spectrophotometry, the removal of pigmented compounds from the lysate prior to CsCl ultracentrifugation could be observed, with absorbance across the visible spectrum decreasing after two washes despite the aqueous volume of the lysate being reduced from 3mL to 1.1mL (Figure 3). Exact pigmented compounds and their partitioning behavior varies between samples, but in all samples isolated, the process corresponds with a reduction in the coloration of the lysate. It is also beneficial to reduce the volume of the crude DNA lysate prior to preparation of the CsCl gradient without precipitation, as precipitation may entrap DNA with contaminating compounds. During the phase separation between a NaCl solution and 2-butoxyethanol, the volume of each phase is dependent on the concentration of NaCl (Table S1). This behavior enables a greater lysis volume and greater sample inputs, increasing the efficiency of the protocol.

**Figure 3.**
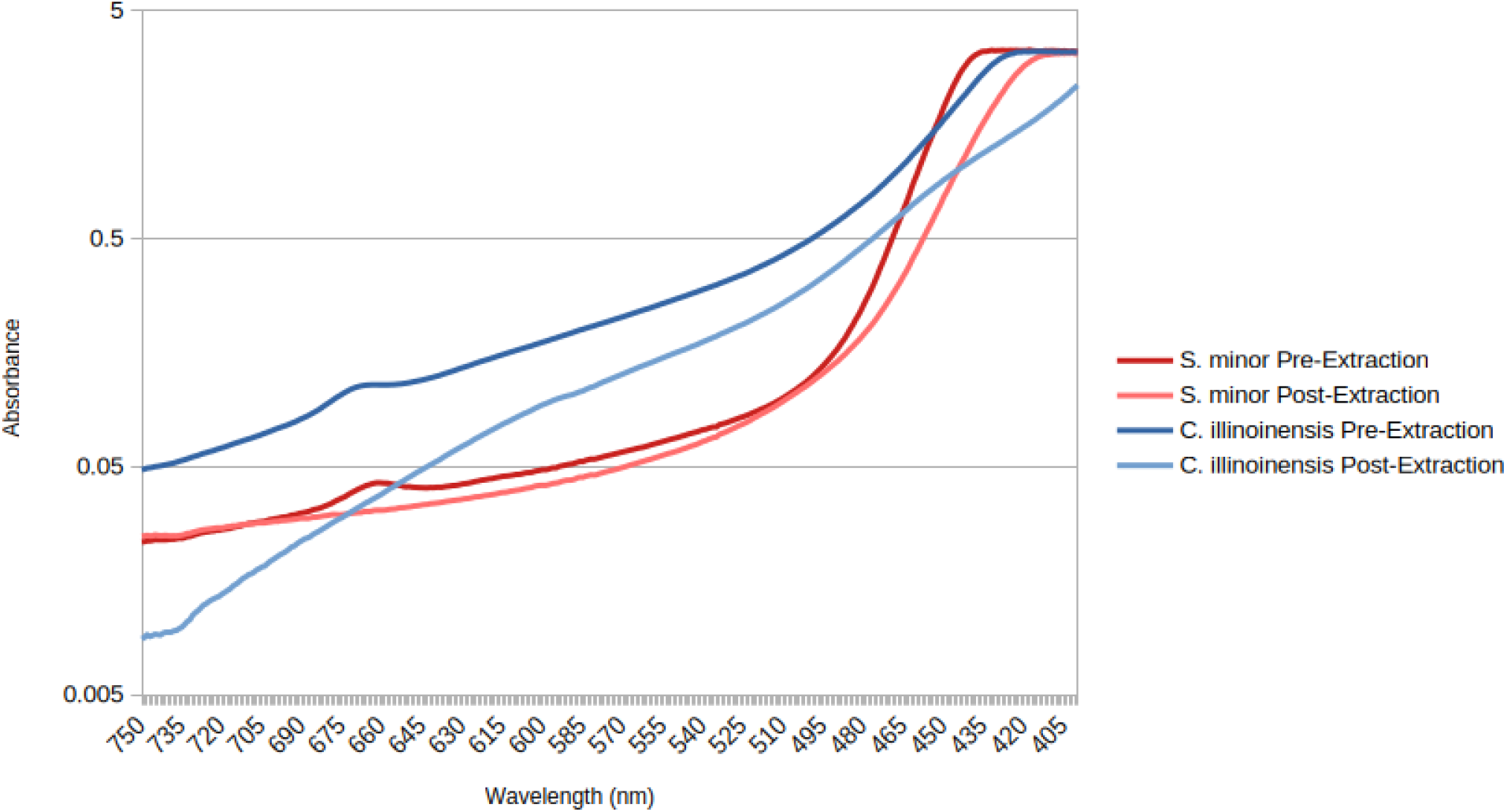
Absorbance of two samples (*Sabal minor* and *Carya illinoinensis*) between 750nm and 400nm before and after two 2-butoxyethanol extractions.

A commonly cited objection to the use of cesium chloride ultracentrifugation is a safety concern in using ethidium bromide [19,32]. With the availability of DNA-specific dyes with reduced toxicity, ethidium bromide can be substituted, however, and dyes such as SYBR Safe and GelGreen have been used previously [33,34]. SYBR Gold was shown to be compatible with a cesium chloride gradient in this protocol, showing strong fluorescence (Figure 4). A limitation to this method is the material and labor cost relative to other methods. Whereas silica column-based methods often require minimal hands-on time (approximately 2 hours for a batch of 12 samples in this work), both the modified CTAB method and the CsCl ultracentrifugation-based method required considerably more hands-on time. Additionally, some steps in the CsCl ultracentrifugation-based method are more technically challenging than in the silica column and modified CTAB-based methods, such as the loading and unloading of an ultracentrifuge. The materials for the modified CTAB-based method were less expensive than those of the other two methods tested here, but the results were relatively poor as well (Tables 1 and 2). Considering the cost of Oxford Nanopore flow cells (1,040 USD per R10.4.1 flow cell as of March 1, 2026), reduced costs from gains in sequencing efficiency utilizing this CsCl ultracentrifugation-based method may outweigh the additional costs, especially where deep sequencing of relatively few samples is required. Sample recalcitrance and experimental design will determine which method is the most advantageous.

**Figure 4.**
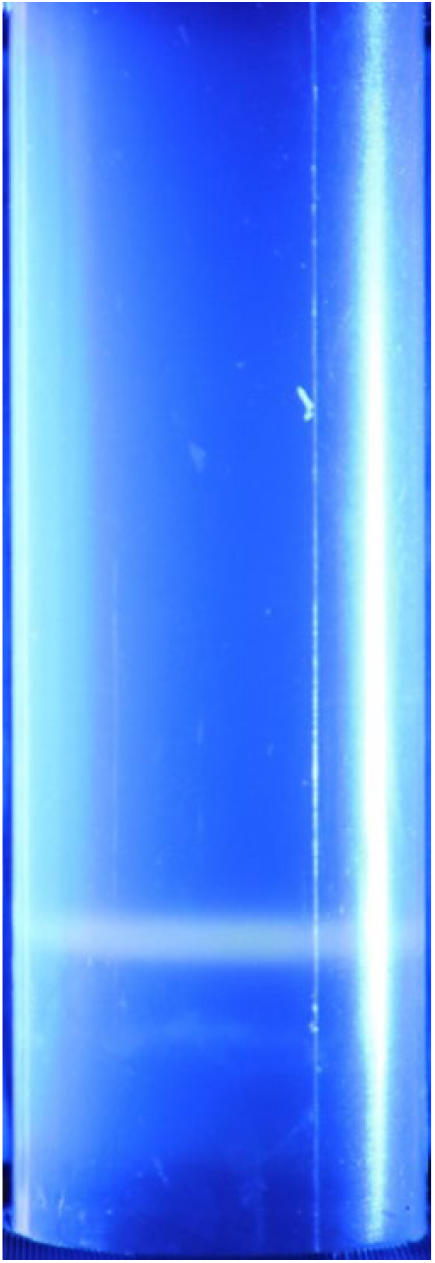
A completed CsCl gradient illuminated with 445nm light just prior to DNA extraction.

## Conclusion

Improved DNA quality and downstream native DNA sequencing success was achieved relative to the most common DNA isolation methods in current use utilizing a modified CsCl ultracentrifugation-based method. This improvement was seen in particularly recalcitrant lineages where standard methods performed poorly, but also in samples relatively amenable to DNA isolation. When applied to a moderately challenging taxon (*Vitis*), DNA isolated via the modified CsCl ultracentrifugation method produced sequencing outputs improved according to 3 key metrics (total data yield, read length, and read quality) relative to a modified CTAB-derived control. The primary drawbacks of this method relative to existing methods are an increase in hands-on time per sample and the need for an ultracentrifuge. As many plant taxa produce difficulties in isolating high-quality, sequencing-ready DNA, however, this method may still be widely applicable. Increased sequencing yield enabled by this method may enable more samples to be sequenced using the same quantity of resources, and greater read N50 and read quality may result in improved genomic assemblies.

## References

1. Koren S, Bao Z, Guarracino A, Ou S, Goodwin S, Jenike KM, et al. Gapless assembly of complete human and plant chromosomes using only nanopore sequencing. Genome Res. Cold Spring Harbor Lab; 2024;34:1919–30.

2. Hu H, Wang J, Nie S, Zhao J, Batley J, Edwards D. Plant pangenomics, current practice and future direction. Agriculture Communications [Internet]. 2024;2:100039. 10.1016/j.agrcom.2024.100039

3. Jayakodi M, Shim H, Mascher M. What Are We Learning from Plant Pangenomes? Annu Rev Plant Biol. Annual Reviews; 2024;76.

4. Labbancz J, Dhingra A. Tree Fruit and Nut Crops at the Dawn of the Pangenomic Era. Horticulturae. Multidisciplinary Digital Publishing Institute (MDPI); 2025. 10.3390/horticulturae11121537

5. Mitchell N, McAssey E V., Hodel RGJ. Emerging methods in botanical DNA/RNA extraction. Appl. Plant Sci. John Wiley and Sons Inc; 2023. 10.1002/aps3.11530

6. Ribeiro RA, Lovato MB. Comparative analysis of different DNA extraction protocols in fresh and herbarium specimens of the genus Dalbergia. Genet Mol Res. 2007;6:173–87.

7. Hadi M, Stacy EA. An optimized RNA extraction method for diverse leaves of Hawaiian Metrosideros, a hypervariable tree species complex. Appl Plant Sci. John Wiley and Sons Inc; 2023;11. 10.1002/aps3.11518

8. Rezadoost MH, Kordrostami M, Kumleh HH. An efficient protocol for isolation of inhibitor-free nucleic acids even from recalcitrant plants. 3 Biotech. Springer Verlag; 2016;6:1–7. 10.1007/s13205-016-0375-0

9. Zhang T, Li H, Jiang M, Hou H, Gao Y, Li Y, et al. Nanopore sequencing: flourishing in its teenage years. Journal of Genetics and Genomics [Internet]. 2024;51:1361–74. 10.1016/j.jgg.2024.09.007

10. Silva C, Machado M, Ferrão J, Sebastião Rodrigues A, Vieira L. Whole human genome 5’-mC methylation analysis using long read nanopore sequencing. Epigenetics. Taylor and Francis Ltd.; 2022;17:1961–75. 10.1080/15592294.2022.2097473

11. Demeke T, Jenkins GR. Influence of DNA extraction methods, PCR inhibitors and quantification methods on real-time PCR assay of biotechnology-derived traits. Anal. Bioanal. Chem. 2010. p. 1977–90. 10.1007/s00216-009-3150-9

12. Schenk JJ, Becklund LE, Carey SJ, Fabre PP. What is the “modified” CTAB protocol? Characterizing modifications to the CTAB DNA extraction protocol. Appl Plant Sci. Wiley Online Library; 2023;11:e11517.

13. Doyle JJ, Doyle JL. A rapid DNA isolation procedure for small quantities of fresh leaf tissue. Phytochemical bulletin. 1987;

14. Mayjonade B, Gouzy J, Donnadieu C, Pouilly N, Marande W, Callot C, et al. Extraction of high-molecular-weight genomic DNA for long-read sequencing of single molecules. Biotechniques. Eaton Publishing Company; 2016;61:203–5. 10.2144/000114460

15. Šimková H, Číhalíková J, Vrána J, Lysák MA, Doležel J. Preparation of HMW DNA from plant nuclei and chromosomes isolated from root tips. Biol Plant. Springer; 2003;46:369–73.

16. Manen JF, Sinitsyna O, Aeschbach L, Markov A V., Sinitsyn A. A fully automatable enzymatic method for DNA extraction from plant tissues. BMC Plant Biol. 2005;5. 10.1186/1471-2229-5-23

17. Richards E, Reichardt M, Rogers S. Preparation of Genomic DNA from Plant Tissue. Curr Protoc Mol Biol [Internet]. John Wiley & Sons, Ltd; 1994;27:2.3.1-2.3.7. 10.1002/0471142727.mb0203s27

18. Green MR, Sambrook J. Preparation of plasmid dna by alkaline lysis with sodium dodecyl sulfate: Maxipreps. Cold Spring Harb Protoc. Cold Spring Harbor Laboratory Press; 2018;2018:pdb-prot093351.

19. Shin JH. Nucleic Acid Extraction Techniques. In: Tang Y-W, Stratton CW, editors. Advanced Techniques in Diagnostic Microbiology [Internet]. Boston, MA: Springer US; 2013. p. 209–25. 10.1007/978-1-4614-3970-7_11

20. Jagielski T, Gawor J, Bakuła Z, Zuchniewicz K, Zak I, Gromadka R. An optimized method for high quality DNA extraction from microalga Prototheca wickerhamii for genome sequencing. Plant Methods. BioMed Central Ltd.; 2017;13. 10.1186/s13007-017-0228-9

21. Firozi P, Zhang W, Chen L, Quiocho FA, Worley KC, Templeton NS. Identification and removal of colanic acid from plasmid DNA preparations: Implications for gene therapy. Gene Ther. 2010;17:1484–99. 10.1038/gt.2010.97

22. Schneider WC, Shelton E, Kuff EL. Association of DNA with melanin granules. J Natl Cancer Inst. Oxford University Press; 1975;55:665–70.

23. Sharma P, Purohit SD. An improved method of DNA isolation from polysaccharide rich leaves of Boswellia serrata Roxb. Indian J. Biotechnol. 2012.

24. Bokszczanin K, Przybyla AA. New simple and efficient method of DNA isolation from pear leaves rich in polyphenolic compounds. Int J Hortic Sci. 2006;12:21–4.

25. Kavidayal H, Rawat A, Saroj S, Ginwal HS. An improved and effective DNA extraction protocol for Pyracantha crenulata with optimal PCR reliability. Silvae Genet. Sciendo; 2024;73:110–9. 10.2478/sg-2024-0011

26. Ou S, Liu J, Chougule KM, Fungtammasan A, Seetharam AS, Stein JC, et al. Effect of sequence depth and length in long-read assembly of the maize inbred NC358. Nat Commun. Nature Research; 2020;11. 10.1038/s41467-020-16037-7

27. Manning K. Isolation of nucleic acids from plants by differential solvent precipitation. Anal Biochem [Internet]. 1991;195:45–50. 10.1016/0003-2697(91)90292-2

28. Ellis CM. The 2-butoxyethanol-water system: Critical solution temperatures and salting-out effects. J Chem Educ [Internet]. American Chemical Society; 1967;44:405. 10.1021/ed044p405

29. Shang Z, Xu P, Feng T, Li X. Insight into the aggregation and phase behavior for aqueous solution of ethylene glycol monobutyl ether with self-assembly induced optical and electrical properties. J Mol Liq [Internet]. 2023;390:123206. 10.1016/j.molliq.2023.123206

30. Bunge AL, Persichetti JM, Payan JP. Explaining skin permeation of 2-butoxyethanol from neat and aqueous solutions. Int J Pharm. 2012;435:50–62. 10.1016/j.ijpharm.2012.01.058

31. Sangster J. Octanol-Water Partition Coefficients of Simple Organic Compounds. J Phys Chem Ref Data [Internet]. 1989;18:1111–229. 10.1063/1.555833

32. Ali N, Rampazzo R de CP, Costa ADT, Krieger MA. Current Nucleic Acid Extraction Methods and Their Implications to Point-of-Care Diagnostics. Biomed Res Int [Internet]. John Wiley & Sons, Ltd; 2017;2017:9306564. 10.1155/2017/9306564

33. Gao J, Pan K, Li H, Fan X, Sun L, Zhang S, et al. Application of GelGreenTM in cesium chloride density gradients for DNA-stable isotope probing experiments. PLoS One. Public Library of Science; 2017;12. 10.1371/journal.pone.0169554

34. Martineau C, Whyte LG, Greer CW. Development of a SYBR safeTM technique for the sensitive detection of DNA in cesium chloride density gradients for stable isotope probing assays. J Microbiol Methods [Internet]. 2008;73:199–202. 10.1016/j.mimet.2008.01.016

